# A lenvatinib-resistance-derived transcriptional program identifies metabolic identity remodeling associated with unfavorable survival in hepatocellular carcinoma

**DOI:** 10.64898/2026.08.21.746217

**Authors:** Li Zheng, Lu Gan

## Abstract

**Background:** Metabolic adaptation is a recognized feature of therapeutic resistance in hepatocellular carcinoma (HCC), but it is unclear whether transcriptional states exposed during acquired resistance are restricted to drug adaptation or reflect broader aggressive tumor biology. We tested whether metabolic programs derived from a lenvatinib-resistance model identify a clinically adverse transcriptional state in an independent HCC patient cohort.

**Methods:** The discovery framework was based on GSE186191, comprising parental and acquired lenvatinib-resistant Hep3B and Huh7 cells. A pre-specified 33-gene lipid-source ledger served as a biological anchor, and three discovery-derived programs—MYC Targets V2, mTORC1 Signaling, and Fatty Acid Metabolism—were frozen before patient-level evaluation. In TCGA-LIHC, single-sample enrichment scores for the three programs were population-standardized and summed to generate an integrated metabolic score. Overall survival was assessed by Kaplan–Meier and Cox analyses. Whole-transcriptome differences between high– and low-score tumors were characterized by preranked gene set enrichment analysis (GSEA).

**Results:** The survival cohort comprised 282 patients (118 deaths), with 141 patients in each median-defined score group. High-score patients had shorter overall survival (log-rank P=0.000419). The continuous score was associated with mortality in univariable analysis (HR 1.86, 95% CI 1.33–2.61; P=0.000293) and in the frozen model adjusted for age, sex, and stage indicators (HR 1.93, 95% CI 1.35–2.76; P=0.000350; n=277). In 327 primary tumors, Fatty Acid Metabolism was strongly depleted in high-score tumors (NES –2.06; FDR<0.001). MYC Targets V2 (NES 1.18; FDR=0.232) and mTORC1 Signaling (NES 1.11; FDR=0.229) showed positive directional enrichment without FDR significance.

**Conclusions:** A lenvatinib-resistance-derived transcriptional program is associated with an adverse-survival state in HCC. The strongest patient-level pathway feature is depletion of canonical fatty-acid metabolism, accompanied by directional MYC/mTORC1 features rather than statistically established pathway activation. These findings support a testable model of metabolic identity remodeling but do not establish causality or clinical prediction of lenvatinib response.

## Introduction

Hepatocellular carcinoma (HCC) is characterized by substantial molecular, metabolic, and phenotypic heterogeneity. Although tumor stage, hepatic reserve, and treatment response remain major determinants of clinical outcome, genomic and transcriptomic studies have shown that HCC tumors also occupy distinct biological states that are not fully captured by conventional clinicopathological variables [1,2]. Metabolic plasticity is a central component of this heterogeneity because malignant hepatocytes must continuously adjust nutrient acquisition, biosynthetic demand, energy production, redox balance, and lipid handling to sustain tumor growth.

This plasticity is particularly relevant under therapeutic stress. Lenvatinib is a multikinase inhibitor used in advanced HCC, yet acquired resistance limits durable disease control. Experimental resistance models show that prolonged drug pressure can induce broad transcriptional and signaling adaptation [3]. Such models are commonly used to identify direct mediators of treatment failure. An alternative possibility is that drug resistance functions as a perturbational system that exposes cellular states already accessible to HCC cells during tumor evolution. Under this interpretation, a program observed in resistant cells need not be a specific clinical marker of lenvatinib response; it may instead represent a broader stress-adapted state associated with aggressive tumor behavior.

Metabolic remodeling is a plausible substrate for such convergence. MYC regulates transcriptional programs required for biomass accumulation, nucleotide synthesis, mitochondrial function, and nutrient utilization [4,5]. mTORC1 integrates growth-factor and nutrient signals to coordinate translation and anabolic metabolism [6]. Lipid metabolism is similarly reorganized in cancer through changes in de novo lipogenesis, extracellular lipid acquisition, lipid storage and mobilization, and mitochondrial fatty-acid oxidation [7]. These processes are interconnected, raising the possibility that malignant progression involves transitions between coordinated metabolic states rather than uniform activation of a single pathway.

We established a frozen analytical framework anchored by 33 genes representing six aspects of lipid-source utilization: de novo lipogenesis, free-fatty-acid uptake, lipoprotein/cholesterol acquisition, lipid storage, lipid mobilization, and mitochondrial fatty-acid oxidation. Analysis of the lenvatinib-resistance dataset GSE186191 prioritized three transcriptional programs—MYC Targets V2, mTORC1 Signaling, and Fatty Acid Metabolism—for outcome-independent validation. The resulting patient score is a three-program transcriptional score rather than a direct additive score of the 33 lipid genes; “Frozen33” denotes the lineage of the discovery framework.

We therefore asked whether a metabolic state exposed during acquired lenvatinib resistance could also be detected across primary HCC tumors and whether this state carried clinical information. Using TCGA-LIHC as an independent patient cohort, we reconstructed the three-program score, evaluated its association with overall survival, and examined the whole-transcriptome context of high– and low-score tumors. The data support a conservative model in which a resistance-derived metabolic state is associated with adverse survival and marked depletion of canonical fatty-acid metabolism, while MYC/mTORC1 findings remain directional rather than statistically definitive.

## Materials and methods

### Study design and datasets

This retrospective computational study followed a discovery-to-validation design. The discovery and validation datasets were analytically separated. Gene-set membership and the composition of the three-program metabolic framework were defined before survival analysis in TCGA-LIHC and were not reselected according to patient outcome. The discovery dataset was GSE186191, comprising parental and acquired lenvatinib-resistant Hep3B and Huh7 HCC cells, with three biological replicates for each condition within each cell line [3]. TCGA-LIHC was used as the independent patient-level validation dataset [2]. No new human tissue was collected.

### Frozen33 discovery lineage

The original biological framework contained a prespecified 33-gene ledger covering six components of lipid-source biology: de novo lipogenesis (6 genes), free-fatty-acid uptake (4 genes), lipoprotein/cholesterol acquisition (4 genes), lipid storage (6 genes), lipid mobilization (5 genes), and mitochondrial fatty-acid oxidation (8 genes). This ledger served as a biological anchor rather than a fitted prognostic signature. Pathway-level analysis of the resistance model prioritized MYC Targets V2, mTORC1 Signaling, and Fatty Acid Metabolism for downstream evaluation. The archived gene-set memberships contained 40, 90, and 61 genes, respectively, and were frozen before TCGA survival analysis. Accordingly, Frozen33 refers to the discovery-program lineage and not to a 33-gene additive patient risk score.

### TCGA-LIHC expression and clinical data

TCGA-LIHC gene-expression and clinical data were harmonized using participant identifiers. Primary tumor samples were identified using TCGA sample-type annotations before patient-level survival merging. The final survival-score cohort contained 282 patients. A separate transcriptomic comparison included 327 primary tumors with available integrated-score classification and whole-transcriptome data. The analysis populations are summarized in Table 1.

**Table 1.** Analysis populations and cohort flow.

| Analysis stage | n | Key definition |
| --- | --- | --- |
| GSE186191 discovery | 12 | Hep3B/Huh7; parental vs lenvatinib-resistant; 3 replicates per condition within each cell line |
| Expression samples mapped | 390 | TCGA-LIHC samples available to the reconstruction workflow before final analysis filtering |
| Transcriptomic high-vs-low comparison | 327 | High score n=167; low score n=160 |
| Survival-score cohort | 282 | 118 deaths; 164 censored; median split 141/141 |
| Adjusted Cox cohort | 277 | 114 deaths; five patients without age excluded |

### Single-sample pathway scoring and integrated score

Single-sample gene-set enrichment analysis was performed for MYC Targets V2, mTORC1 Signaling, and Fatty Acid Metabolism using the frozen GMT definitions [8]. The reproducible scoring implementation used rank normalization, weight 0.25, minimum gene-set size 5, maximum gene-set size 500, and no phenotype permutations. In the survival cohort, each pathway score was population-standardized as z=(x-mu)/sigma using the cohort mean and population standard deviation. The integrated metabolic score was the sum of the three standardized component scores. Patients were dichotomized at the cohort median for Kaplan– Meier visualization, whereas the continuous score was used in Cox regression.

### Overall-survival analysis

Overall survival was the clinical endpoint. Kaplan–Meier curves compared median-defined high– and low-score groups using a two-sided log-rank test [9]. The integrated score was also analyzed as a continuous covariate using Cox proportional-hazards regression [10]. The univariable model included 282 patients. The frozen multivariable model included integrated score, age, sex, Stage II, and Stage III–IV indicator variables. Five patients without age information were excluded, yielding n=277. In the archived stage encoding, missing or unclassified stage records shared the all-zero indicator pattern of the Stage I reference category; therefore, the adjusted model is interpreted as a covariate-adjusted association rather than definitive proof of stage-independent prognostic utility.

### Differential-expression and preranked GSEA

For transcriptomic characterization, 327 primary tumors were classified by integrated score (high n=167; low n=160). Gene-level differences were evaluated on log-transformed expression data using a non-parametric Mann–Whitney framework with false-discovery-rate correction. Genes were ranked for preranked GSEA by log2FC × [-log10(P)], with positive values indicating expression toward the high-score phenotype. GSEA used the three frozen programs, weight=1, 1,000 permutations, random seed=42, minimum gene-set size 5, and maximum size 500 [11–13]. MYC Targets V2, mTORC1 Signaling, and Fatty Acid Metabolism contributed 40/40, 89/90, and 61/61 genes, respectively, to the ranked dataset. Directional findings that did not pass FDR correction were treated as trends rather than statistically established activation.

### Reproducibility

The downstream TCGA validation workflow—including single-sample scoring, integrated-score reconstruction, survival analysis, Cox regression, high-versus-low expression comparison, and preranked GSEA—was reconstructed from archived expression, clinical, sample-mapping, score, ranked-list, and GMT files. The maximum absolute difference between the reconstructed and frozen integrated score was below 3×10^-15. Executable code and software-environment records reproducing the TCGA validation workflow were archived with the analytical package.

## Results

### A resistance-derived framework links experimental adaptation to patient-level metabolic states

We used acquired lenvatinib resistance as a perturbational discovery context rather than as a direct surrogate for clinical treatment response. GSE186191 contains parental and acquired lenvatinib-resistant states in Hep3B and Huh7 cells. The Frozen33 lineage was anchored by a 33-gene lipid-source ledger and subsequently represented at pathway level by three frozen programs: MYC Targets V2, mTORC1 Signaling, and Fatty Acid Metabolism (Figure 1). This design allowed a focused test of whether metabolic organization exposed under sustained drug pressure could also be detected in primary human HCC.

**Figure 1.**
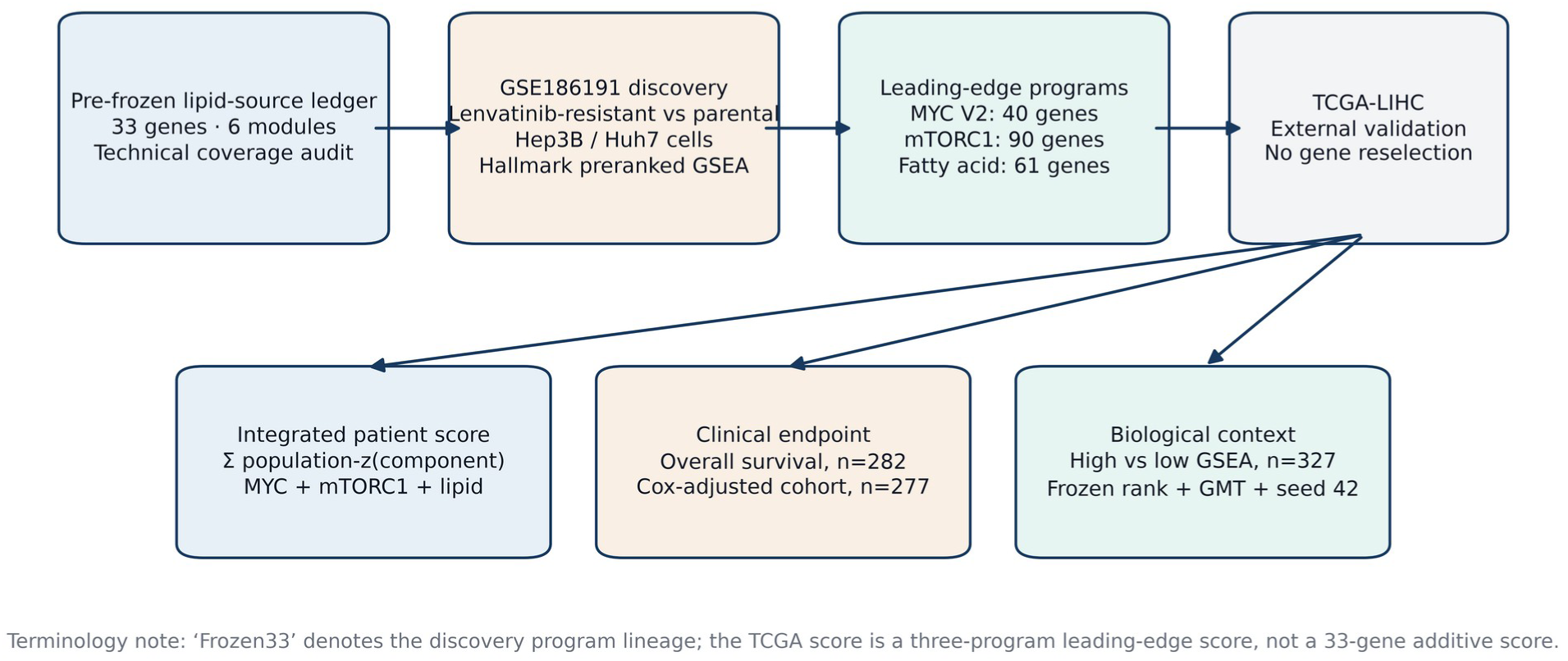
Discovery-to-validation architecture. A pre-frozen 33-gene lipid-source ledger anchored analysis of parental and acquired lenvatinib-resistant Hep3B/Huh7 cells in GSE186191. Discovery-derived transcriptional programs were frozen before external evaluation in TCGA-LIHC. Frozen33 denotes the discovery-program lineage; the TCGA integrated score is a three-program score and not a direct 33-gene additive score.

### The three frozen programs define a continuous metabolic-score landscape in TCGA-LIHC

Single-sample enrichment analysis demonstrated substantial intertumoral variation in the three programs. After population standardization, the component scores were summed to generate the integrated metabolic score. Reconstruction reproduced the archived patient-level score to floating-point precision. Among 282 patients with survival information, the score had a median of –0.0126 and separated the cohort into 141 high-score and 141 low-score patients (Figure 2). The continuous distribution is consistent with a graded tumor state rather than requiring a rigid binary subtype.

**Figure 2.**
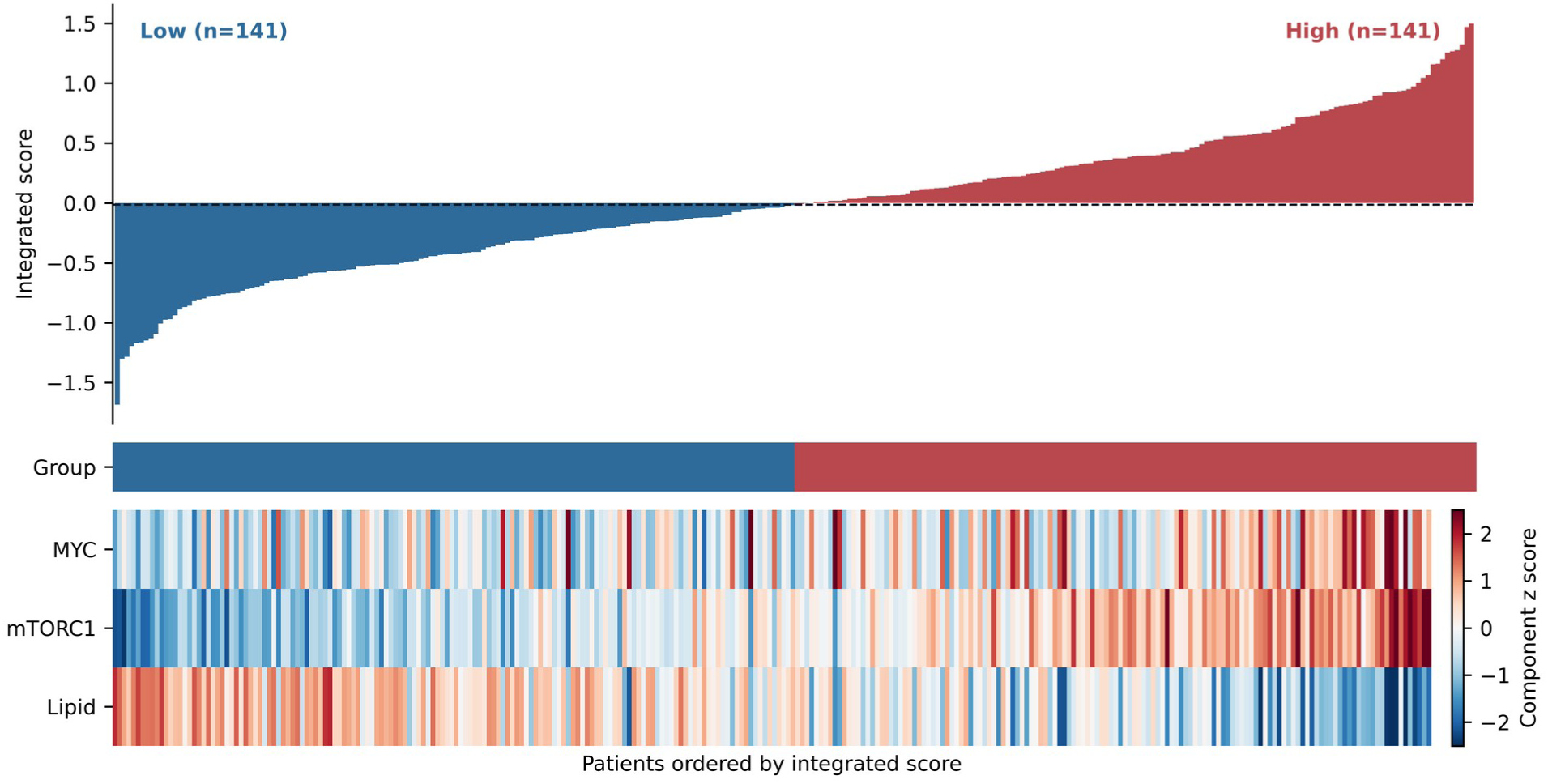
Integrated metabolic-score landscape in the survival cohort. Patients (n=282) are ordered by integrated score. The score is the sum of population-standardized MYC Targets V2, mTORC1 Signaling, and Fatty Acid Metabolism single-sample enrichment scores. Median stratification yielded 141 high-score and 141 low-score patients. The component heatmap shows the standardized score for each program.

### A higher metabolic score is associated with unfavorable overall survival

High-score patients had significantly shorter overall survival than low-score patients (two-sided log-rank P=0.0004189; Figure 3A). As a continuous variable, the score was associated with higher mortality in univariable Cox regression (HR 1.862, 95% CI 1.330–2.607; P=0.000293). The association remained in the frozen model adjusted for age, sex, and stage indicators (HR 1.928, 95% CI 1.345–2.762; P=0.000350; n=277; Figure 3B). The survival estimates are summarized in Table 2. Because TCGA-LIHC was not selected according to lenvatinib exposure or response, these findings identify a survival-associated tumor state and do not establish a clinical predictor of lenvatinib benefit or resistance.

**Figure 3.**
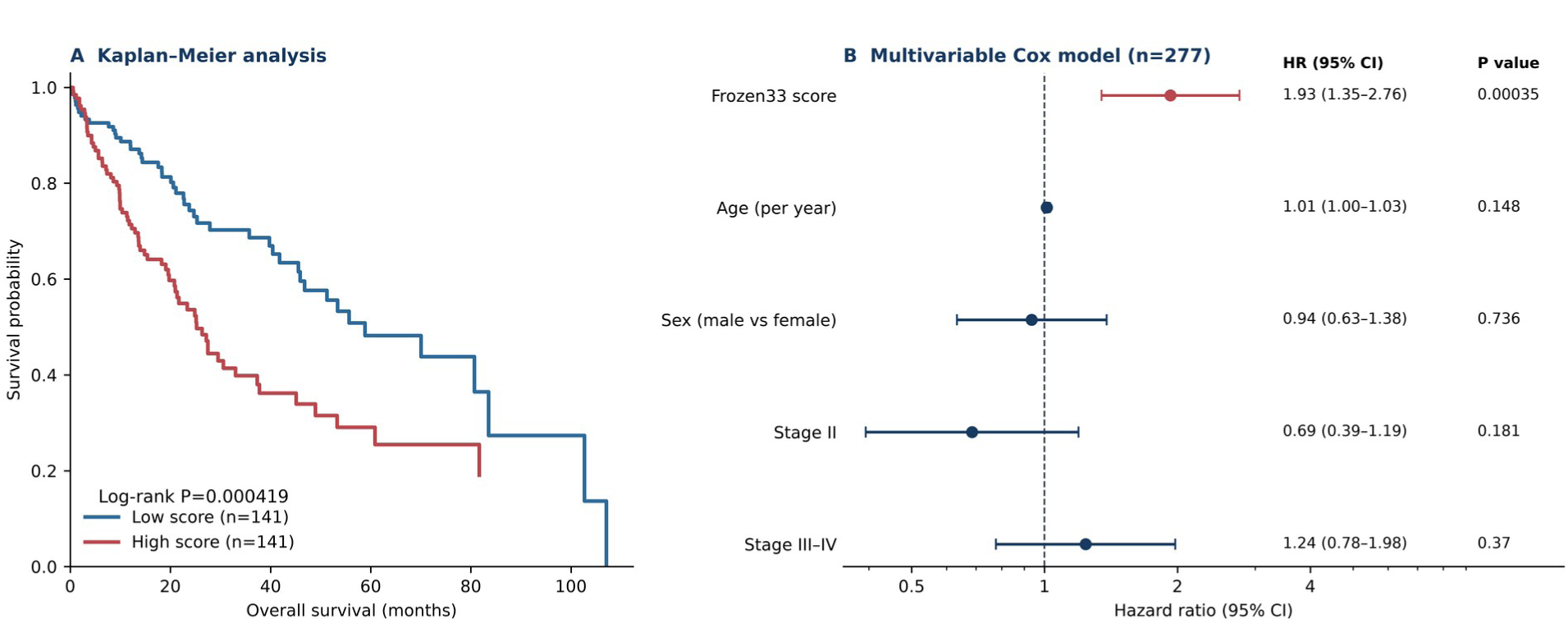
Survival association of the integrated metabolic score. (A) Kaplan–Meier curves compare median-defined score groups (log-rank P=0.000419). (B) Forest plot of the frozen multivariable Cox model (n=277; 114 events), including continuous score, age, sex, Stage II, and Stage III–IV indicators.

**Table 2.** Association of the integrated metabolic score with overall survival.

| Analysis | Estimate | 95% CI | P value | n / events |
| --- | --- | --- | --- | --- |
| Kaplan–Meier log-rank, high vs low | — | — | 0.000419 | 282 / 118 |
| Univariable Cox, continuous score | HR 1.862 | 1.330–2.607 | 0.000293 | 282 / 118 |
| Multivariable Cox, continuous score | HR 1.928 | 1.345–2.762 | 0.000350 | 277 / 114 |

### High-score tumors show marked depletion of canonical fatty-acid metabolism

Whole-transcriptome comparison of 167 high-score and 160 low-score primary tumors identified broad expression differences. Several proliferation-associated genes, including MYBL2, PARPBP, CDC6, KIFC1, RAD51AP1, HJURP, FOXM1, PLK1, GINS1, and CDK1, were increased toward the high-score phenotype. Preranked GSEA showed strong negative enrichment of Fatty Acid Metabolism in high-score tumors (ES –0.711; NES –2.065; nominal P<0.001; FDR<0.001; Figure 4). MYC Targets V2 showed positive enrichment (ES 0.650; NES 1.175; nominal P=0.137; FDR=0.232), and mTORC1 Signaling showed a similar positive direction (ES 0.595; NES 1.115; nominal P=0.191; FDR=0.229). The complete frozen GSEA estimates are summarized in Table 3. Thus, the strongest statistically supported pathway feature of the high-score state was depletion of the canonical fatty-acid metabolic program. MYC and mTORC1 are interpreted as directional contextual features rather than established pathway-activation events.

**Figure 4.**
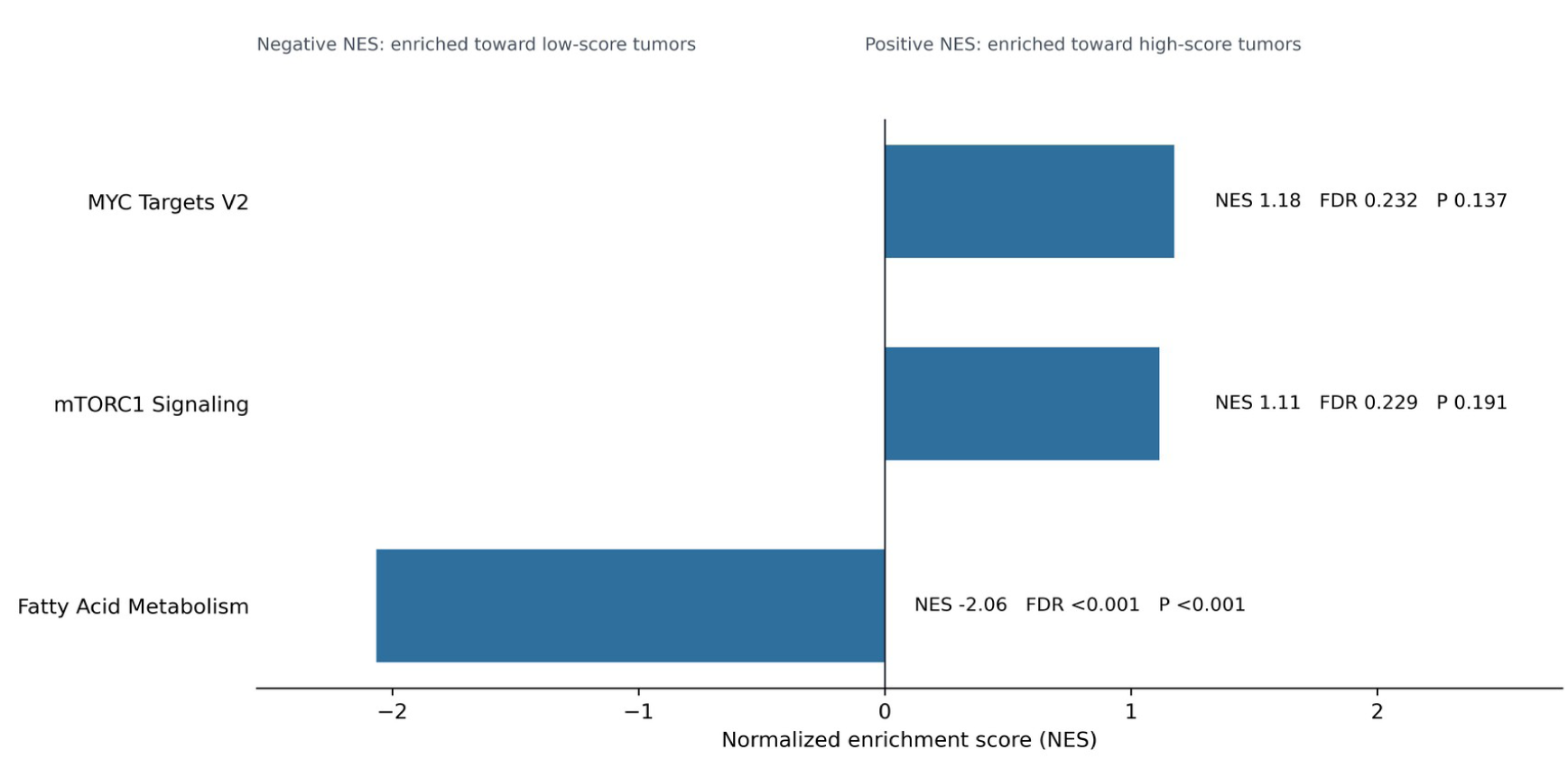
Preranked GSEA of high-versus low-score HCC tumors. In 327 primary tumors, Fatty Acid Metabolism was significantly enriched toward the low-score phenotype (NES –2.065; FDR<0.001). MYC Targets V2 (NES 1.175; FDR=0.232) and mTORC1 Signaling (NES 1.115; FDR=0.229) were directionally enriched toward high-score tumors but did not reach FDR significance.

**Table 3.**
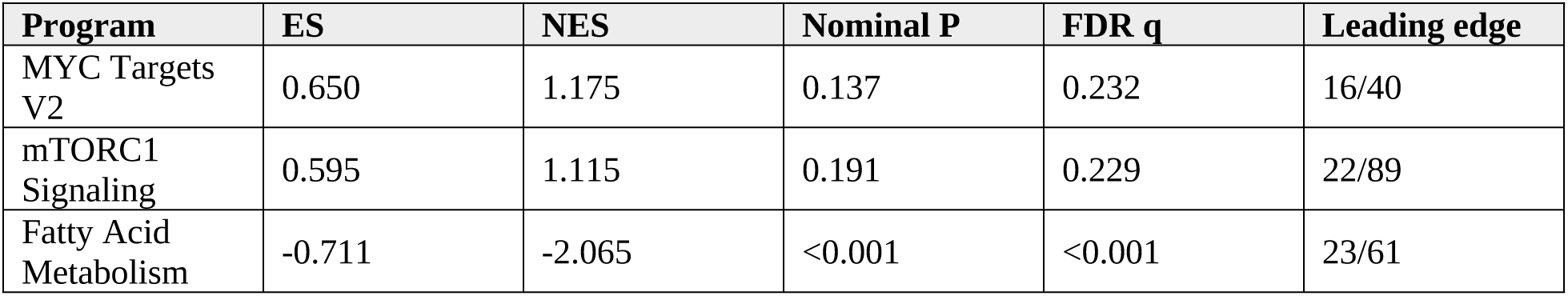
Preranked GSEA of high-versus low-score tumors. Interpretation: Positive NES indicates enrichment toward the high-score phenotype; negative NES indicates enrichment toward the low-score phenotype. Only Fatty Acid Metabolism reached FDR significance.

| Program | ES | NES | Nominal P | FDR q | Leading edge |
| --- | --- | --- | --- | --- | --- |
| MYC Targets V2 | 0.650 | 1.175 | 0.137 | 0.232 | 16/40 |
| mTORC1 Signaling | 0.595 | 1.115 | 0.191 | 0.229 | 22/89 |
| Fatty Acid Metabolism | -0.711 | -2.065 | <0.001 | <0.001 | 23/61 |

### The integrated evidence is consistent with metabolic identity remodeling

The survival and transcriptomic findings together describe a resistance-derived state associated with adverse outcome, increased proliferation-related transcription, marked loss of canonical fatty-acid metabolism, and directional MYC/mTORC1 features (Figure 5). We use the term metabolic identity remodeling to describe this coordinated transcriptomic pattern. The term represents a testable biological interpretation and does not imply that MYC, mTORC1, or altered lipid metabolism has been experimentally shown to cause lenvatinib resistance or adverse survival in the present study.

**Figure 5.**
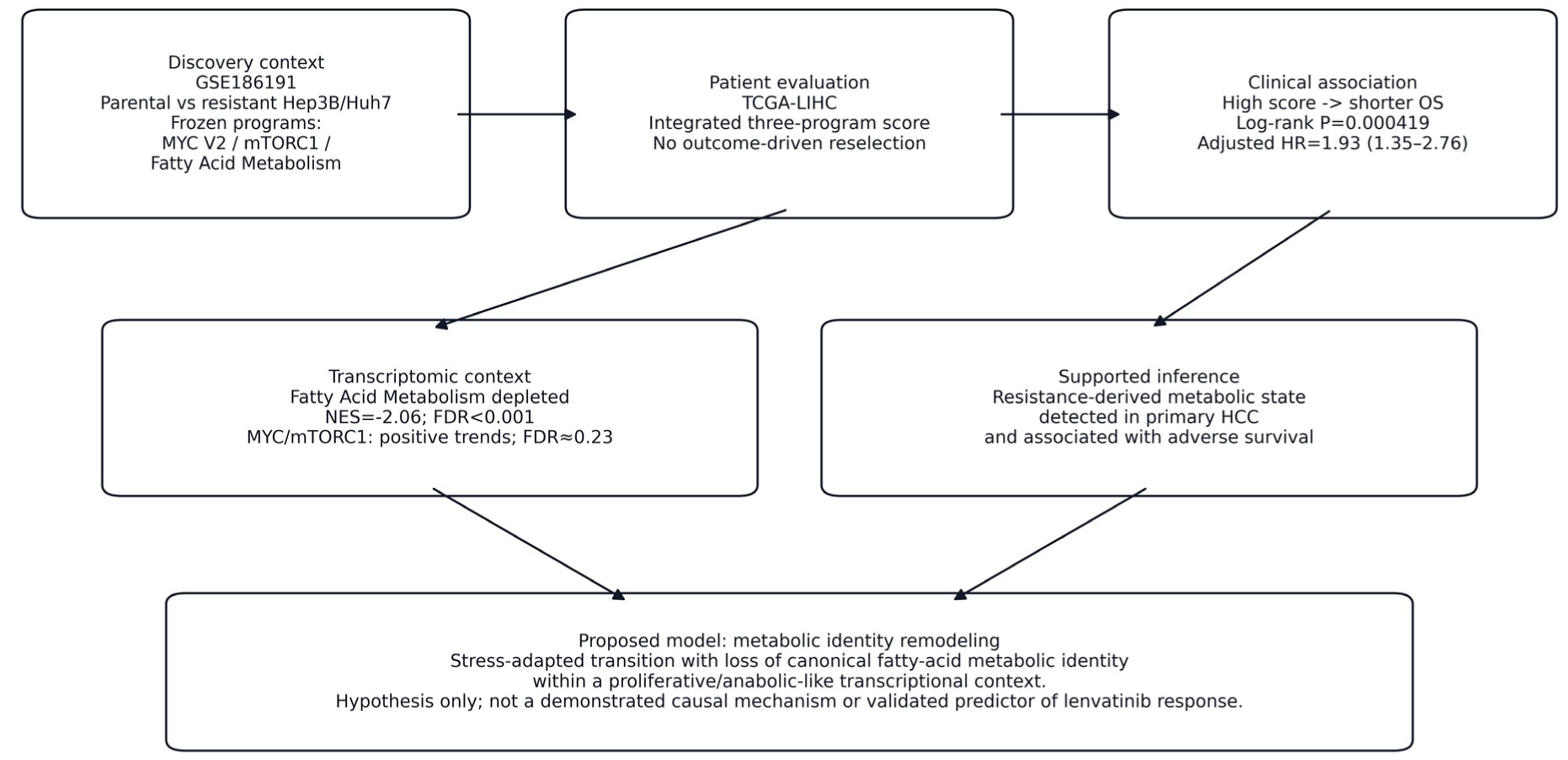
Evidence-supported model of metabolic identity remodeling. Acquired lenvatinib resistance is used as a perturbational discovery context. A related three-program metabolic state is detectable in TCGA-LIHC and is associated with adverse overall survival. The strongest patient-level pathway feature is depletion of canonical Fatty Acid Metabolism, whereas MYC and mTORC1 enrichment remain directional trends. Metabolic identity remodeling is presented as a testable biological model; no causal mechanism or validated prediction of lenvatinib response is claimed.

## Discussion

This study connects an acquired lenvatinib-resistance model with an independent patient cohort and identifies a metabolically defined HCC state associated with adverse overall survival. The central observation is not that the score predicts clinical lenvatinib resistance. Rather, a transcriptional organization exposed under sustained therapeutic stress is also detectable across primary tumors not selected on lenvatinib response and carries survival information.

This distinction separates a drug-resistance biomarker from a drug-resistance-derived tumor state. A treatment-predictive biomarker should be evaluated in patients receiving the relevant therapy and, ideally, demonstrate a treatment-by-biomarker interaction. TCGA-LIHC does not provide that design. Here, GSE186191 is better viewed as a perturbational discovery system: chronic drug pressure forces malignant cells to adapt and can reveal states that may also arise during tumor evolution independently of drug exposure. The presence of a related state in primary HCC and its association with survival are consistent with partial convergence between therapeutic adaptation and aggressive tumor biology.

The metabolic pattern is more nuanced than generalized lipid activation. The most robust patient-level pathway result was depletion of Fatty Acid Metabolism in high-score tumors. Normal hepatocytes are highly metabolically specialized, and hepatocarcinogenesis is accompanied by progressive loss or reorganization of differentiated hepatic functions. Reduced canonical fatty-acid metabolic transcription may therefore reflect a shift away from a differentiated hepatocyte-like metabolic identity rather than a uniform decrease in every lipid flux. The present bulk-transcriptome data cannot distinguish changes in pathway activity from changes in cellular composition or substrate use, and functional metabolic measurements will be required to resolve this point.

At the same time, the high-score transcriptome contained multiple proliferation-associated genes, including MYBL2, FOXM1, PLK1, CDK1, and CDC6. MYC Targets V2 and mTORC1 Signaling were directionally enriched toward high-score tumors, but neither passed FDR correction. The current data therefore do not justify a headline claim that coordinated MYC/mTORC1 activation mechanistically drives adverse outcome. A more defensible interpretation is that the high-score state occurs in a proliferative and anabolic-like transcriptional context, while the best-resolved metabolic feature is loss of the canonical fatty-acid program.

These findings motivate the concept of metabolic identity remodeling. In this model, HCC cells can transition between coordinated metabolic states during tumor evolution or sustained stress. A relatively differentiated lipid-utilization state may give way to a stress-adapted phenotype characterized by reduced canonical fatty-acid metabolic identity, altered substrate handling, increased proliferative demand, and reorganization of nutrient-responsive transcriptional programs. This model is a mechanistic hypothesis generated by the present data, not a demonstrated causal pathway.

The framework generates specific experimental predictions. Parental and lenvatinib-resistant HCC models can be compared directly for fatty-acid uptake, lipid storage, mitochondrial fatty-acid oxidation, de novo lipogenesis, and extracellular lipid dependence. MYC or mTORC1 perturbation can then test whether these pathways contribute to establishment or maintenance of the remodeled state. Patient cohorts with documented lenvatinib exposure will be required to determine whether the state is solely prognostic or also has treatment-predictive relevance.

The distinction between the original 33-gene framework and the validated patient score is also important. Frozen33 originated as a biological ledger describing six dimensions of lipid-source utilization; it was not fitted as a 33-gene survival model. Broader transcriptional programs derived from the discovery framework were frozen before TCGA outcome analysis. This design preserves a discovery-to-validation separation and reduces outcome-driven gene selection.

Several limitations require emphasis. First, the discovery stage is based on two established HCC cell lines, and in-vitro resistance does not reproduce the stromal, vascular, immune, and pharmacological complexity of clinical resistance. Second, TCGA-LIHC was not designed to evaluate lenvatinib response, so the present results establish biological overlap and prognostic association rather than treatment prediction. Third, the frozen multivariable model retained missing-stage records in the reference-indicator pattern, limiting claims of stage independence. Fourth, bulk-tumor transcriptomes reflect malignant-cell state as well as tumor purity and non-malignant cellular composition. Fifth, MYC and mTORC1 enrichment was directional but FDR-nonsignificant. Finally, no functional perturbation was performed, so causal relationships among lenvatinib adaptation, MYC, mTORC1, fatty-acid metabolism, and survival cannot be inferred.

Despite these limitations, the study provides a reproducible observation that metabolic organization identified under acquired lenvatinib resistance is detectable in primary HCC and is associated with adverse survival. The patient-level data refine the original hypothesis by indicating that the dominant pathway feature is loss of canonical fatty-acid metabolic identity rather than generalized lipid-pathway activation. This evidence supports metabolic identity remodeling as a focused framework for subsequent functional testing rather than as a completed causal mechanism.

## Conclusion

A lenvatinib-resistance-derived transcriptional program is associated with unfavorable overall survival in HCC. High-score tumors show strong depletion of the Fatty Acid Metabolism program, a proliferation-associated transcriptomic background, and directional MYC/mTORC1 enrichment. These observations support a testable model of metabolic identity remodeling, while the current evidence establishes association rather than causality or clinical prediction of lenvatinib response. Independent clinical validation and functional experiments are required to determine whether this state represents a therapeutically actionable vulnerability.

## Data availability

The discovery RNA-sequencing dataset is publicly available through the NCBI Gene Expression Omnibus under accession GSE186191. Validation transcriptomic and clinical data were obtained from TCGA-LIHC. Analysis-ready expression matrices, sample mappings, clinical tables, patient-level scores, ranked gene lists, frozen GMT definitions, figure source data, and reproducibility records are archived with the analytical package.

## Code availability

Executable Python code and a software-environment record reproducing the downstream TCGA validation workflow are archived with the analytical package. The workflow reconstructs patient-level single-sample enrichment scores, the integrated metabolic score, Kaplan–Meier and Cox analyses, differential-expression ranking, preranked GSEA, and the principal validation figures.

## Ethics statement

This study used publicly available, de-identified cell-line and patient datasets. No new patient specimens were collected and no identifiable participant information was accessed.

## Generative AI assistance

The authors used OpenAI’s ChatGPT (version GPT-5.6, accessed August 2026) solely to assist with language editing and improve the readability of the manuscript. All scientific judgments, interpretations, and conclusions were independently made by the authors. The authors take full responsibility for all scientific content of the manuscript.

## Author contributions

Li Zheng: Conceptualization; Methodology; Formal analysis; Data curation; Visualization; Writing – original draft. Lu Gan: Conceptualization; Writing – review & editing; Supervision. All authors reviewed and approved the manuscript.

## Supporting information

GSEA reproduction files

